# Autism genes are selectively targeted by environmental pollutants including pesticides, heavy metals, bisphenol A, phthalates and many others in food, cosmetics or household products

**DOI:** 10.1101/062521

**Authors:** C.J. Carter, R.A. Blizard

**Author notes:** Correspondence: PolygenicPathways, Flat 2, 40 Baldslow Road, Hastings, East Sussex, TN34 2EY, UK.

## Abstract

The increasing incidence of autism suggests a major environmental influence. Epidemiology has implicated many candidates and genetics many susceptibility genes. Gene/environment interactions in autism were analysed using 206 autism genes (ASG’s) to interrogate ~1 million chemical/gene interactions in the comparative toxicogenomics database. Bias towards ASG’s was statistically determined for each chemical. Many suspect compounds identified in epidemiology, including tetrachlorodibenzodioxin, pesticides, particulate matter, benzo(a)pyrene, heavy metals, valproate, acetaminophen, SSRI’s, cocaine, bisphenol A, phthalates, polyhalogenated biphenyls, flame retardants, diesel constituents, terbutaline and oxytocin, *inter alia* showed a significant degree of bias towards ASG’s, as did relevant endogenous agents (retinoids, sex steroids, thyroxine, melatonin, folate, dopamine, serotonin). Numerous other endocrine disruptors selectively targeted ASG’s including paraquat, atrazine and other pesticides not yet studied in autism and many compounds used in food, cosmetics or household products, including tretinoin, soy phytoestrogens, aspartame, titanium dioxide and sodium fluoride. Autism polymorphisms are known to influence sensitivity to some of these chemicals and these same genes play an important role in barrier function and control of respiratory cilia sweeping particulate matter from the airways. The close gene/environment relationships, for multiple suspect pollutants, suggest that the rising incidence of autism might be chemically driven by numerous environmental contaminants in a gene dependent manner. The protective dappled camouflage of the peppered moth was rendered invalid by industrial soot covering the trees, a situation reversed by clean air acts. The rising tide of neurodevelopmental and other childhood disorders linked to multiple pollutants may need a similar solution.

## Introduction

According to the Center for disease control (CDC) http://www.cdc.gov/ncbddd/autism/data.html the USA incidence of autism spectrum disorders rose 2.2 fold from 2000 to 2010 [1]. In the UK, a five-fold increase in autism in the 1990’s, reached a plateau in the 2000’s up to 2010 [2]. This increased prevalence is likely partly due to environmental influences, of which there are many candidates. Many chemical classes or specific chemicals related to autism have been reviewed by Rossignol and co-authors [3]. These include pesticides (chlorpyrifos, dicofol dialkylphosphate,endosulfan, the Dichlorodiphenyltrichloroethane (DDT) metabolite o,p′-dichlorodiphenyltrichloroethane, polychlorinated dibenzo-p-dioxin, polychlorinated and Polybrominated diphenyls, dichlorodiphenyldichloroethylene, heavy metals (aluminium, arsenic, cadmium, mercury, lead, nickel and manganese), air pollutants (carbon monoxide, diesel components, methylene chloride, nitrogen dioxide, ozone, particulate matter, quinoline, styrene, sulphur dioxide, trichloroethylene and vinyl chloride) and other pollutants (xylene), bisphenol A and phthalates.

Other classes or named pollutants and drugs implicated in autism include 2,3,7,8-tetrachlorodibenzo-p-dioxin ( a contaminant of the Agent orange defoliant) [4], organochlorine pesticides (chlordan) [5], organophosphates and pyrethroids [6], and metals (high hair levels of aluminium, arsenic, cadmium, mercury, antimony, nickel, lead, and vanadium [7] or high urine levels of aluminium, tin, tungsten or thallium [8, 9]. Living in areas with higher air levels of styrene and chromium during pregnancy has also been associated with increased autism risk [10]. Zinc deficiency, excess copper levels, and low Zn/Cu ratio are common in children diagnosed with autism [11] and low levels of iron have been reported [12]. Autism has also been linked to flame retardants (polybrominated diphenyl ether-28 or trans-nonachlor) [5],air pollutants (benzo(a)pyrene [13],nitric oxide [14], 1,3-butadiene, meta/para-xylene, other aromatic solvents, lead, perchloroethylene, and formaldehyde [15], Diethyl phthalate, Di-butylphthalate and di-(2-ethylhexyl) phthalate metabolites ((5-OH-MEHP [mono-(2-ethyl-5-hydroxyhexyl) 1,2-benzenedicarboxylate] and 5-oxo-MEHP [mono-(2-ethyl-5-oxohexyl) 1,2-benzenedicarboxylate) [16, 17]. Heavy smoking [18], or the use of certain drugs during pregnancy, including alcohol and cocaine [18, 19], methamphetamine (case report[20]), acetaminophen [21, 22], valproate [23] as well as thalidomide [24], or selective serotonin uptake inhibitors (SSRI’s) (fluoxetine, sertraline, paroxetine, citalopram, and escitalopram) [25] have also been linked to the development of autism in the offspring, as has the use of pharmaceutical agents for the induction or delay of labour (oxytocin, prostaglandins [26], dexamethasone [27] or terbutaline [28]). Smoking, cannabis use, nicotine and alcohol have also been linked to autistic behaviour in young adults [29]. Autism related genes are preferentially expressed prenatally in the frontal cortex suggesting that an inherent genetic susceptibility may be confined to this period [30].

A number of compounds detailed above have been shown to produce autism-relevant behavioural effects in laboratory models when administered prenatally. For example endocrine disruptors (atrazine, perfluorooctanoic acid, bisphenol-A, 2,3,7,8-tetrachlorodibenzo-p-dioxin) alone or combined in a mixture, from gestational day 7 until weaning produce disrupted behaviour, which for mixture effects was predominantly seen in male mice offspring [31]. Other endocrine disrupting mixtures, (di-n-butylphthalate, diethylhexylphthalate, vinclozolin, prochloraz, procymidone, linuron, epoxiconazole, and DDE) or (bisphenol A, 4-methylbenzylidene camphor, 2-ethylhexyl 4-methoxycinnamate, and butylparaben), when administered prenatally to rats have been shown to modify the expression of genes related to glutamatergic function, the migration and pathfinding of GABAergic and glutamatergic neurones and of autism-related genes in the offspring [32]. The organophosphate insecticide, chlorpyrifos, also induces relevant behavioural effects in mice offspring when administered during pregnancy, again showing male preference [33, 34]. Gestational exposure to heavy metals (cadmium, lead, arsenate, manganese, or mercury) or valproic acid also produces multiple behavioural abnormalities that persist into adulthood in male mice offspring, effects that are accompanied by epigenetic changes in gene methylation [35]

Genes associated with autism are catalogued at the Autworks database using a confidence score derived from analysis of the Genotator association database [36, 37]. 206 genes are regarded as prime autism susceptibility candidates and these genes and network analyses are available at the autworks site from the Wall lab at Harvard University http://tools.autworks.hms.harvard.edu/gene_sets/580/genes.

This same set of genes has recently been shown to be localised and enriched in many barriers including the blood brain barrier, as well as skin, intestinal, placental and trophoblast barriers. Several also play an important role in relation to respiratory cilia that sweep noxious particles from the airways. These barrier-related genes are thus in a position to modify the access of numerous environmental agents to the blood and brain and their role in respiratory cilia is relevant to particulate matter and airborne pollutants [38].

Given the strength of the various environmental associations with autism, and its increasing prevalence over recent years, it is possible that the environmental influences that target these genes may afford clues as to the combined and conditional causes of autism.

Chemical influences on the 206 Autworks susceptibility genes (ASG’s) were analysed using the Comparative Toxicogenomics Database (CTD) [39] which records over 1 million interactions between diverse chemicals and genes or proteins. Previous work using this database has already shown a link between autism or other disease-related genes and environmental risk factors [40]. For example, asthma has been linked with p,p’-DDT, and autism with o,p’-DDT, both metabolites of the organochlorine insecticide dichlorodiphenyltrichloroethane (DDT) [41].

The results show that the toxicogenomic effects of many chemicals associated with autism selectively target the ASG’s, showing a close relationship between genes and environment.

## Methods

206 Autworks autism susceptibility genes (ASG’s) http://tools.autworks.hms.harvard.edu/gene_sets/580/genes [36] were analysed. Gene definitions are provided in supplementary File 1. Members of this gene set are highlighted in bold when they appear in the text. The gene symbols (applicable to human genes and many mouse or rat homologues) were uploaded to the Comparative Toxicogenomics Database (CTD) [39] http://ctdbase.org/. All interactions are referenced at CTD and can be accessed by uploading the gene symbols from the Autworks dataset. The results were downloaded and the number of ASG’s and the total number of genes (autism and others affected by each chemical or the number of chemicals affecting each autism gene were curated manually. Chemicals were broadly classified into groups (e.g. pesticides, metals, endocrine disruptors). Singletons (chemicals affecting only one gene) were ignored. Many clinical and research drugs were returned, but are not treated in this paper.

All chemicals possess a unique CAS registry number, from the American Chemical society Chemical Abstracts Service http://www.cas.org/content/chemical-substances allowing cross-referencing between CTD data and compounds in other databases. Overlaps were identified using the Venny tool http://bioinfogp.cnb.csic.es/tools/venny/ [42].

The databases used for such classification, based largely on overlapping CAS numbers, included The TEDX List of Potential Endocrine Disruptors http://endocrinedisruption.org/; The EU list of endocrine disruptors http://eng.mst.dk/topics/chemicals/endocrine-disruptors/the-eu-list-of-potential-endocrine-disruptors/, the NIST Polycyclic Aromatic Hydrocarbon (PAH) Structure Index http://pah.nist.gov/,the national toxicity program from the US department of health http://ntp.niehs.nih.gov/index.cfm and the United States Environmental protection agency databases http://www.epa.gov/. Persistent organic pollutants (POPs) are as defined by the Stockholm convention http://chm.pops.int/Home/tabid/2121/mctl/ViewDetails/EventModID/871/EventID/514/xmid/6921/Default.aspx.

Compounds in cigarettes are defined by the Federal drug administration http://www.fda.gov/TobaccoProducts/GuidanceComplianceRegulatoryInformation/ucm297786.htm and the Tobacco Products Scientific Advisory Committee list of harmful or potentially harmful components in tobacco and/or tobacco smoke [43].

Compounds found in diesel exhaust are listed at Wikipedia http://en.wikipedia.org/wiki/Dieselexhaust and at the United States department of labor Partial List of Chemicals Associated with Diesel Exhaust https://www.osha.gov/SLTC/dieselexhaust/chemical.html

Lists of chemicals in cosmetics, foods and pharmaceutical preparations were obtained from the National Research Council (US) Steering Committee on Identification of Toxic and Potentially Toxic Chemicals for Consideration by the National Toxicology Program [44], the UK Food standards agency listing EU approved food additives http://www.food.gov.uk/ and from the International fragrance association http://www.ifraorg.org/en/ingredients#.U_w5JWNWpZx. Food ingredients were also interrogated at FooDB http://foodb.ca/compounds a project from the Canadian Metabolomics Innovation Centre.․ Food additives are also listed at GSFA online http://www.codexalimentarius.net/gsfaonline/additives/index.html from the Joint FAO/WHO Expert Committee on Food Additives (JECFA) and from the List of Indirect Additives Used in Food Contact Substances from the US Food and drug administration http://www.accessdata.fda.gov/scripts/fcn/fcnNavigation.cfm?rpt=iaListin and the EAFUS list (Everything Added to Food in the United States) http://www.accessdata.fda.gov/scripts/fcn/fcnNavigation.cfm?rpt=eafusListing&;displayAll=true.

The Consumer Product Information Database (CPID) http://whatsinproducts.com/contents/aboutcpid/1 was use for ingredients found in household products. Chemicals in household products were identified using the National Institute of health Household Products database http://householdproducts.nlm.nih.gov/cgibin/household/search.

It is important to appreciate that the only selection criteria were the 206 autism genes which were sent to forage chemical interactions in an extensive toxicogenomics database, and that the compounds returned are unbiased by any other factor.

## Gene enrichment analysis

The ASG’s, selected by Autworks by confidence score based on Genotator, number 206 (0.77% from a human genome of 26,846 protein-coding genes). There were 10,766 unique chemicals in CTD, with 1,002,333 curated interactions (2015 data). If a chemical affects N genes, one would expect an equal proportion of ASG’s (0.77%) to be contained within this gene set (Expected = N*(206/26846)). Chemical bias towards the ASG’s is reflected by observed/expected ratios >1 and the corresponding p value derived from the hypergeometric probability test, which was corrected for false discovery [45], with a final cut-off at P<0.05. Most results are illustrated graphically. For individual compounds the data are illustrated by N autism genes affected/total number of genes affected by the compound, followed by the fold enrichment and p values (e.g. Methionine (69/3724: 2.41 fold: P= 2.E-12).

## Results

67861 chemical/gene interactions affected the ASG’s. 4428 compounds affected 1 or more ASG’s. 6338 chemicals did not interact with any autism gene. The number of ASG’s targeted by each chemical varied from 1 to 141 (Tetrachlorodibenzodioxin). The number of chemicals affecting each autism gene ranged from 0 to 1669. 760 compounds with significant enrichment values affected ≥5 ASG’s; 372 ≥10; 109 ≥25; 29 ≥50; 6 ≥100. Enrichment values (observed/expected ASG’s per total 9number of genes affected by each compound) for these significant chemicals, where the number of ASG’s targeted > 5 ranged from 1.4 to 97.7. No chemical interactions had been curated for **HTR3C, KLF14, RP1L1** or **ZNF778**

### Genes affected by compounds implicated in autism (Fig 1)

Of the named pollutants implicated in autism (see introduction) 43 showed enrichment values at P < 0.05 (all except Nickel, diethyl phthalate, Nitrogen Dioxide and Vinyl Chloride) Compounds with the most significant enrichment scores were pesticides (diazinon, chlorpyrifos, Dichlorodiphenyldichloroethylene (DDE: a DDT metabolite) and cypermethrin), and metals (arsenic, zinc, mercury and cadmium) Other highly significant pollutants included nitric oxide, Bisphenol A, benzo(a)pyrene, particulate matter and ozone (Fig 1).

**Figure 1.**
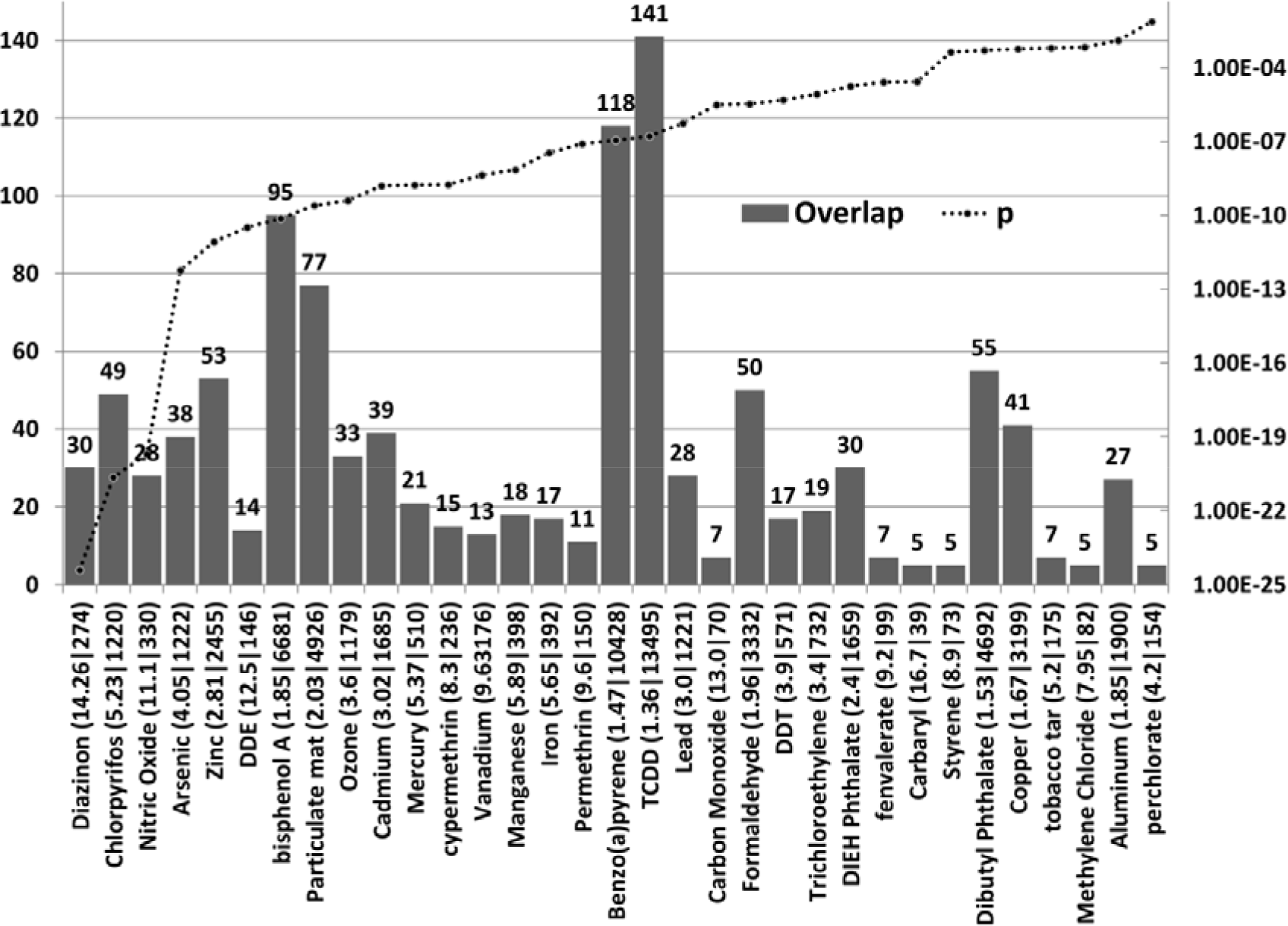
The number of ASG’s (where N>=5) affected by pesticides, herbicides, heavy metals and other named pollutants implicated in autism (left axis). The enrichment ratio and the total number of genes affected by each compound are shown after each compound name. For example Diazinon affects 274 genes in total, 30 of which are ASG’s, yielding an enrichment value (observed/expected) of 14.26. (Diazinon (14.26|274). FDR corrected p values are shown on the right hand Y axis, which is set at a maximum of p=0.05. TCDD= Tetrachlorodibenzodioxin. DDT = dichlorodiphenyltrichloroethane: DDE= Dichlorodiphenyl Dichloroethylene (DDT metabolite). DIEH Phthalate = Diethylhexyl Phthalate Dicofol, Sulphur Dioxide, Chlordan, acephate, cyhalothrin, quinoline, 3-xylene and 1,3-butadiene overlaps were also significant but affected less than 5 ASG’s (not shown).

Drugs with the most significant enrichment scores included SSRI antidepressants (fluoxetine, sertraline, paroxetine, citalopram); thalidomide, drugs of abuse (cocaine, methamphetamine, and ethanol); nicotine, steroid drugs (dexamethasone and hydrocortisone) and drugs used to induce (Dinoprost, misoprost, oxytocin) or prevent (terbutaline) labour in pregnancy, as well as thalidomide, acetaminophen, thimerosal and valproate (Fig 2). There have been multiple conflicting studies relating to the risks and benefits of Thimerosal containing vaccines, which have resulted in its withdrawal in many countries [46]. Thimerosal was removed from childhood vaccines in the USA in 2001. http://www.cdc.gov/vaccinesafety/Concerns/thimerosal/index.html.

Thimerosal affects 8 autism genes (**GSTM1, IL6, MAPK1, MAPK3, PTK2, RFC1, SLC1A1** and **TNF**) and its relatively minor enrichment effects (compared to many industrial and other pollutants) may well be limited to those with particular polymorphisms in this set. Such gene dependence is also seen in different strains of mice. Thimerosal administration mimicking childhood immunization is not toxic in C57BL/6J or BALB/cJ mice. Autoimmune disease-sensitive SJL/J mice show growth delay, hypolocomotion, exaggerated response to novelty and densely packed, hyperchromic hippocampal neurons with altered glutamate receptors and transporters [47].

Together, these results show that a large number of industrial, agrochemical and household pollutants or drugs implicated in autism selectively target multiple ASG’s. One evident interpretation is that polymorphisms therein may modify sensitivity to autism-related chemicals. This is discussed in a later section. Using a similar experimental approach Kauchik et al constructed a protein/protein interaction (PPI) network of autism related genes and found that the effects of drug mixtures (environmental contaminant concentrations of carbamazepine, venflaxine and fluoxetine) or clinical concentrations of valproate on gene expression in fish brains or in human neuronal cell cultures tended to target the same networks as those identified in the autism PPI interactome [48].

**Figure 2.**
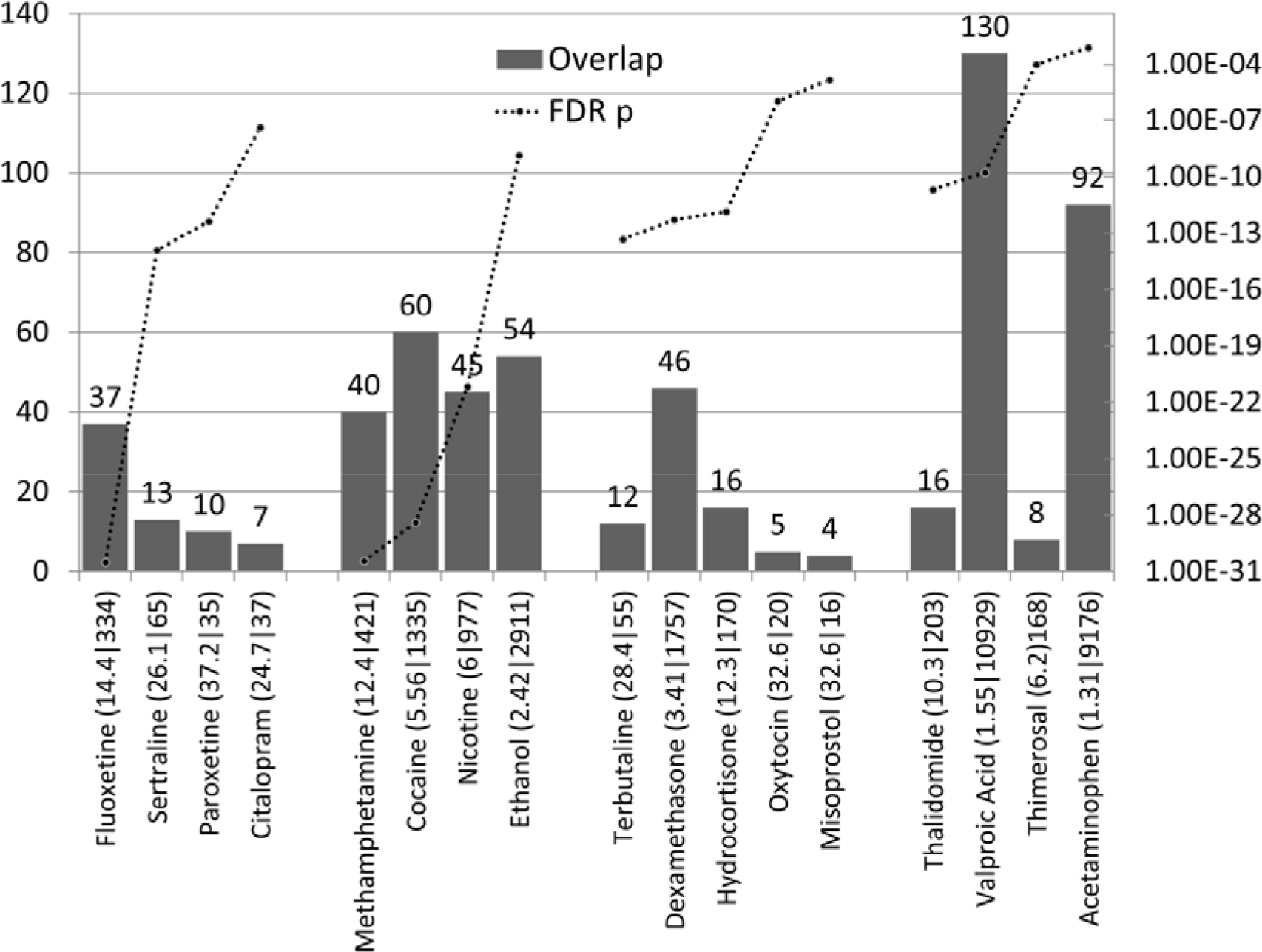
The number of ASG’s affected by drugs implicated in autism (left axis), and p values (right axis). The enrichment ratio and the total number of genes affected by each compound are shown after each compound name. First batch = SSRI antidepressants, second = drugs of abuse, third = drugs used during labour, fourth =others.

### Other pesticides, fungicides and herbicides

Many pesticides, other than those reportedly related to autism (see Fig 1), are used agriculturally or in the home, often together or at different seasons. 41 of these targeted multiple autism genes (P< 0.05) (Fig 3). In the various classes, Cycloheximide, Maneb, Vinclozolin and mancozeb were the most significant fungicides; Paraquat, Cyanazine,Atrazine and herbimycin the most significant herbicides and Methoxychlor, Dieldrin, Endosulfan, chlorpyrifos oxon the highest scoring insecticides.

**Figure 3.**
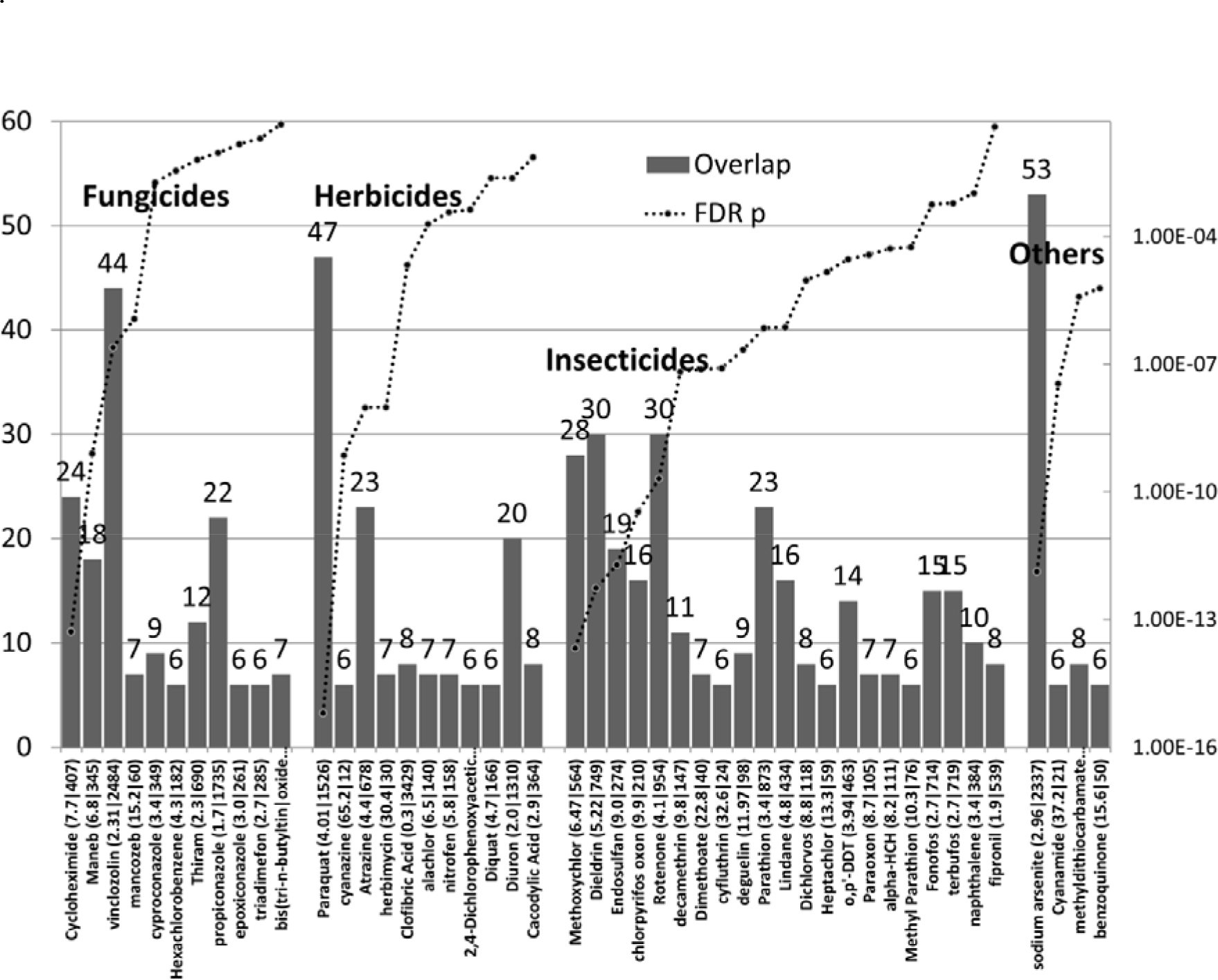
The number of ASG’s affected by diverse pesticides (left axis), and p values (right axis). The enrichment ratio and the total number of genes affected by each compound are shown after each compound name. The compounds are divided by class: Others includes diverse broad-spectrum pesticides and Cyanimide, which is widely used in agriculture to promote uniform opening of buds, early foliation and bloom in fruits.

### Other metals

For the most part, other metals significantly orienting their effects towards the ASG’s were salts of those already shown in Fig 1 (arsenic, zinc, cadmium, mercury, lead, copper, aluminium) (not shown).). Asbestos, Crocidolite (28/682: 5.4 fold: p= 2.1E-12) is blue asbestos, a product linked to many cancers but not studied in relation to autism. The metals also included the highly toxic tributyltin (10/228: 5.7 fold: p=2.45E-05), a suspected carcinogen, cobaltous chloride (49/3281: 1.95 fold: p= 4.92E-06. Titanium dioxide (50/3449:1.8 fold: p=8.3E-06) and silicon dioxide (37/2789: 1.7 fold: p=0.0006) are included in the EAFUS and cosmetics lists and treated in these sections.

### Poly-halogenated biphenyls, flame retardants, bisphenols and phthalates

4 known flame retardants (all polybromo biphenyls) and 12 polychlorinated biphenyls showed significant enrichment values in relation to the ASG’s as did several bisphenols and phthalates (Fig 4)

**Figure 4.**
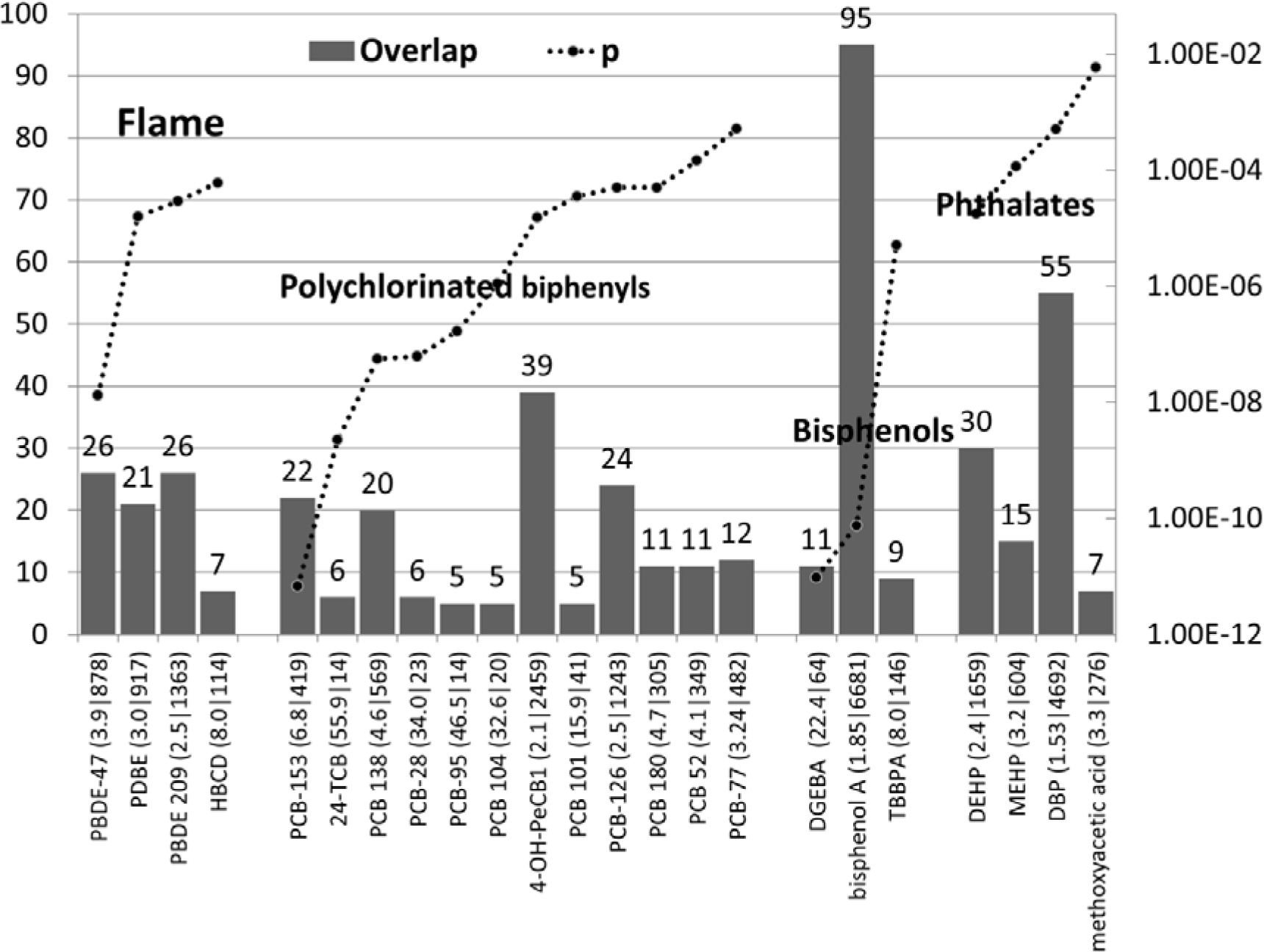
The number of ASG’s affected by diverse polyhalogenated biphenyls, bisphenols and phthalates (left axis) and p values (right axis). The enrichment ratio and the total number of genes affected by each compound are shown after each compound name. Flame= Flame retardants. Methoxyacetic acid is a di(2-methoxyethyl) phthalate metabolite. PBDE-47 =2,2′,4,4′-tetrabromodiphenyl ether; PDBE =pentabromodiphenyl ether; PBDE 209 =decabromobiphenyl ether; HBCD hexabromocyclododecane; PCB-153 =2,4,5,2′,4′,5′-hexachlorobiphenyl; 24-TCB =2,4,2′,4′-tetrachlorobiphenyl; PCB 138 =2,2′,3′,4,4′,5-hexachlorobiphenyl; PCB-28 =2,4,4′-trichlorobiphenyl; PCB-95 =2,2′,3,5′,6-pentachlorobiphenyl; PCB 104 =2,2′,4,6,6′-pentachlorobiphenyl; 4-OH-PeCB1 =2′,3,3′,4′,5-pentachloro-4-hydroxybiphenyl; PCB 101 =2,4,5,2′,5′-pentachlorobiphenyl; PCB-126 =3,4,5,3′,4′-pentachlorobiphenyl; PCB 180 =2,2′,3,4,4′,5,5′-heptachlorobiphenyl; PCB 52 =2,5,2′,5′-tetrachlorobiphenyl; PCB-77 =3,4,3′,4′-tetrachlorobiphenyl; DGEBA =bisphenol A diglycidyl ether; TBBPA =tetrabromobisphenol A; DEHP =Diethylhexyl Phthalate; MEHP =mono-(2-ethylhexyl|phthalate; DBP =Dibutyl Phthalate

### Persistent organic pollutants (POP) and Polycyclic aromatic hydrocarbons (PAH)

Several of these compounds, already recognised for their toxicity in many domains significantly targeted the ASG’s (Fig 5). A large number of genes were targeted by 2,3,7,8-Tetrachlorodibenzodioxin and Benzo(a)pyrene.

**Figure 5.**
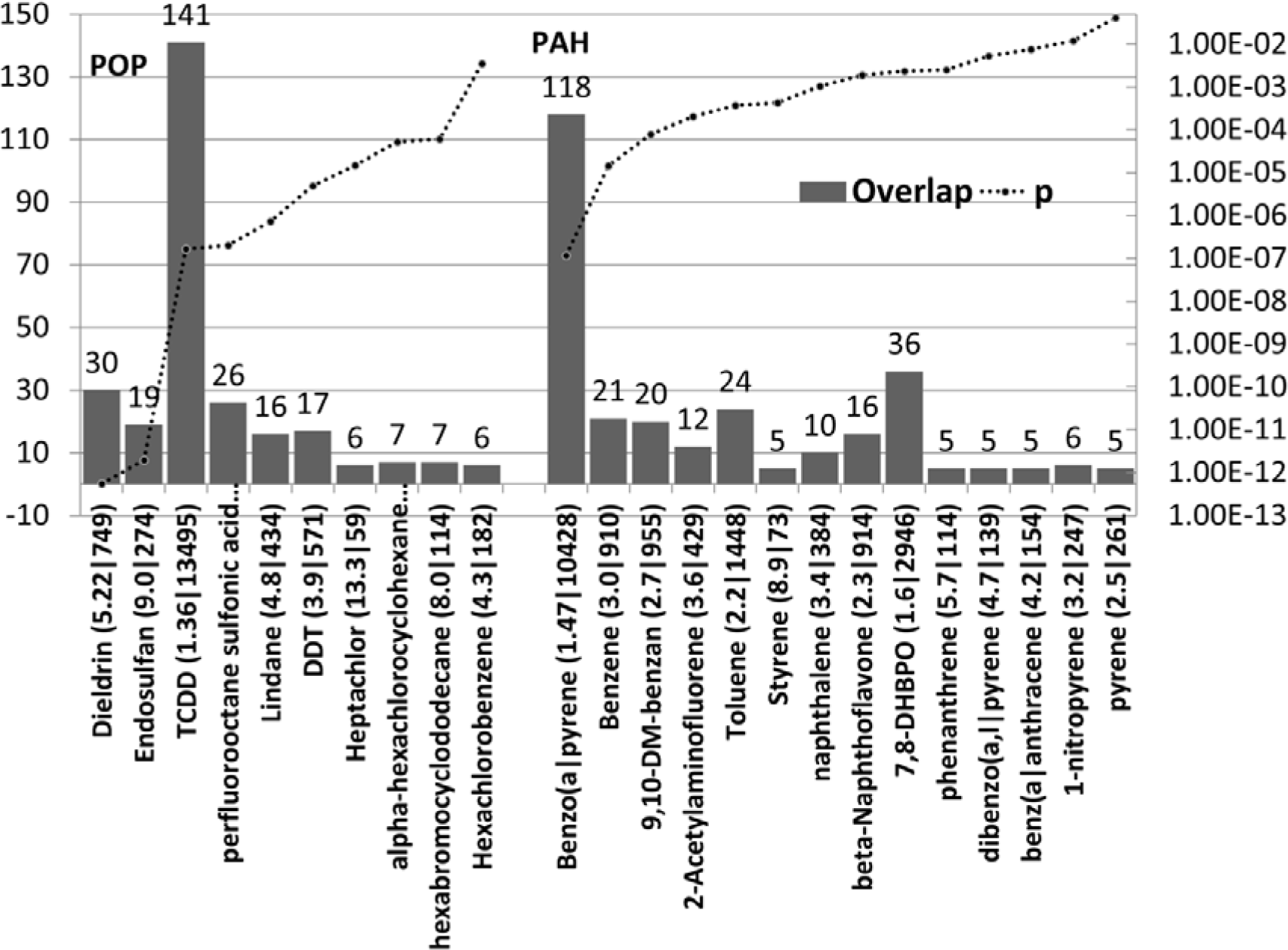
The number of ASG’s affected by diverse Persistent organic pollutants (POP) and Polycyclic aromatic hydrocarbons (PAH). (left axis), and p values (right axis). The enrichment ratio and the total number of genes affected by each compound are shown after each compound name. DDT =dichlorodiphenyltrichloroethane; TCDD = 2,3,7,8-Tetrachlorodibenzodioxin; 9,10-DM-benzan = 9,10-Dimethyl-1,2-benzanthracene; 7,8-DHBPO = 7,8-Dihydro-7,8-dihydroxybenzo(a)pyrene 9,10-oxide.

### Endocrine disruptors

138 compounds within this class significantly oriented their effects towards 5 or more autism genes, 79 > 10 genes, 39 >20 genes, 16 > 30 genes (P<0.05). Many of these compounds are in the EAFUS list or in food, as plant constituents (e.g. phytoestrogens) or contaminants (e.g. alkylphenols). Many are also found in cosmetics or household products and these classes are coded for in Fig 6. Several of these compounds, for example pesticides, heavy metals, bisphenols and phthalates, have already been treated above and are not included in Fig 6. The highly significant endocrine disruptors also include several used in cosmetics including bisphenol A, nonylphenol, Butylated Hydroxytoluene, Butylated Hydroxyanisole, beta Carotene, 4-propylphenol, Styrene, acetyl methyl tetramethyl tetralin, 4-cresol, 4-ethylphenol, Resorcinol, Phenol, Ethylene Glycol, Propylparaben and n-hexane.

Several plant-derived phytoestrogens, flavones/flavonoids, selectively target these genes (Apigenin, daidzein, Genistein, kaempferol, Luteolin, naringenin, resveratrol and quercetin). They are common components of food supplements, including baby milk, follow-ons, and soy formula [49–51]. They are generally regarded as potentially beneficial in a number of conditions including cancer, type 2 diabetes, obesity, coronary heart disease, metabolic syndrome, and neurodegenerative diseases. (e.g. resveratrol [52]). Luteolin and quercetin have been reported to reduce autism symptoms in a small clinical trial [53]. However, such compounds are not bereft of toxicological effects. For example neonatally administered genistein in mice later reduces female fertility and embryo implantation [54]. It is also embryotoxic in rats and synergises with Bisphenol A in this respect [55, 56]. Pre- or perinatally administered phytoestrogens can also have deleterious effects on animal behaviour. For example adult male mice perinatally exposed to daidzein show significantly less exploration and higher levels of anxiety and aggression [57]. Genestein given to rat dams during late pregnancy and early lactation affects the differentiation of brain structures as well as changes in anxiety and aggressive behaviour in the male offspring [58].In Phytoestrogens can be found in pregnant women’s serum and amniotic fluid during pregnancy and soy ingestion increases amniotic fluid phytoestrogen concentrations in female and male foetuses [59].

These endocrine disruptors also include Sodium Fluoride, which is added to domestic water supplies for dental health in many countries. NaF decreases fertility in female rats, via decreases in serum estradiol and progesterone levels and the uterine expression of the follicle stimulating hormone receptor. It also increases uterine estrogen receptor alpha (ESR1) and progesterone receptor and luteinising hormone receptor protein expression levels (400;401). In mouse Leydig tumor cells NaF decrease the mRNA expression of steroidogenic acute regulatory protein (STAR) and a cytochrome P450 (CYP11A1) which catalyses the conversion of cholesterol to pregnenolone, the first rate-limiting step in the synthesis of steroid hormones (402). When given to pregnant rats, NaF decreases the activity levels of testicular steroidogenic marker enzymes (3beta hydroxysteroid dehydrogenase and 17beta hydroxysteroid dehydrogenase) in the 90 day old male offspring (403). Dietary NaF also decreases the serum levels of free and bound triodothronine and thyroxine in rats (404) NaF also decreases the expression of CYP1A2 in mouse spermatozoa (405). CYP1A2 metabolises polycyclic aromatic hydrocarbons, dioxins, polychlorinated dibenzofurans, polychlorinated biphenyls, and acetaminophen (406). NaF thus possesses endocrine disrupting properties and an ability to affect the metabolism of a number of environmental agents implicated in autism.

**Figure 6.**
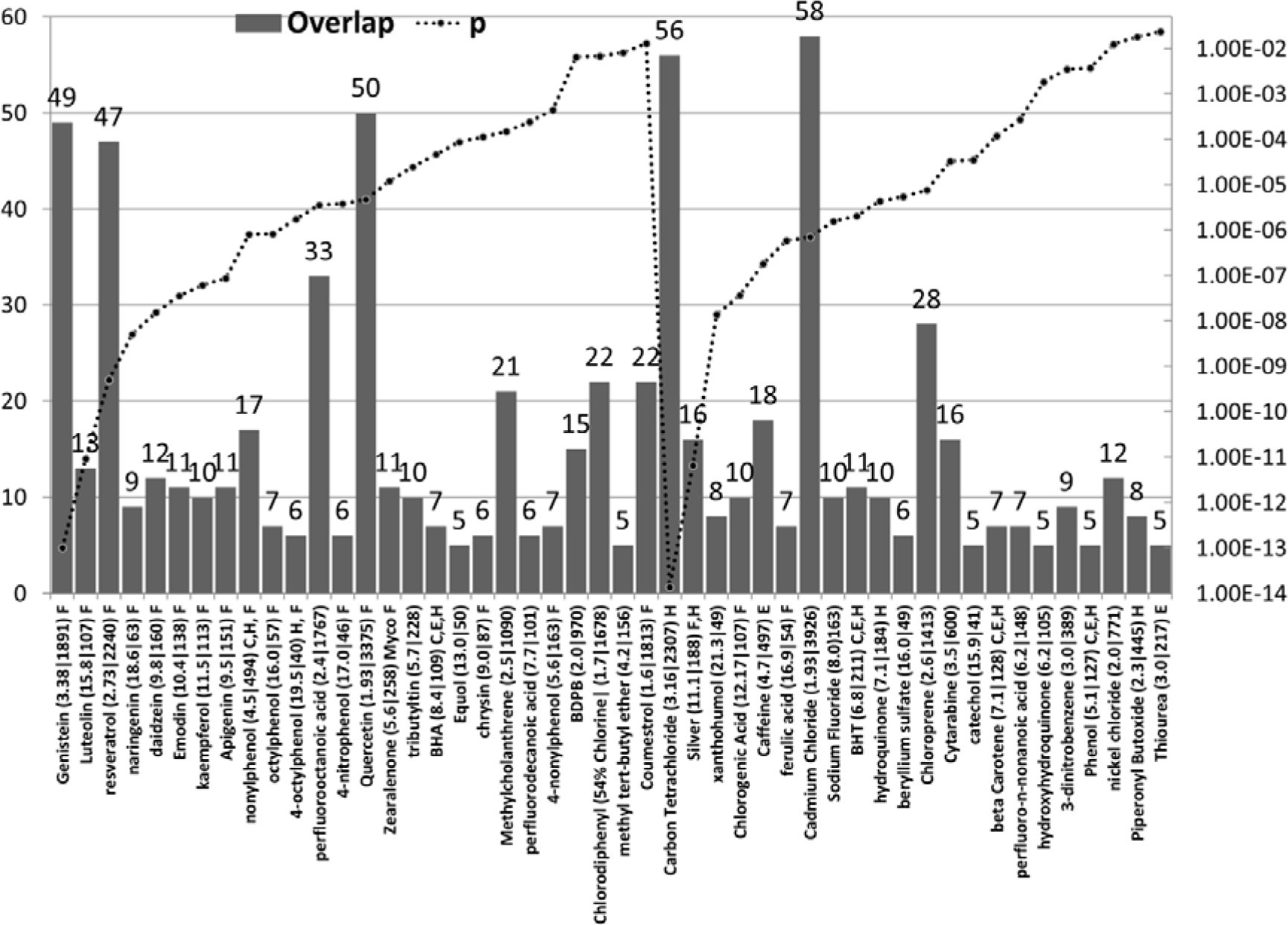
The number of ASG’s affected by diverse known (first batch) and potential (second batch) endocrine disruptors. N ASG’s =left axis, p values =right axis). The enrichment ratio and the total number of genes affected by each compound are shown after each compound name. Also appended are compounds found in Food (plant constituents or contaminants (F), the EAFUS list of food additive (E), Cosmetics (C) and household objects (H). Myco = mycotoxin; BDPB = 1,4-bis(2-(3,5-dichloropyridyloxy))benzene; BHA = Butylated Hydroxyanisole; BHT= Butylated Hydroxytoluene;

### Components of cigarette smoke or diesel exhaust

Many chemicals found in diesel and/or cigarette smoke significantly targeted a number of ASG’s (Fig 7). Their effects must be considered as cumulative.

These data suggest an important relationship between the cumulative effects of smoking or diesel toxicants and ASG’s. In relation to diesel and traffic pollution, a recent review has highlighted air pollution as a contributory factor to both neurodevelopmental and adult neurodegenerative disorders. Air pollution causes oxidative stress and neuro-inflammation, and brain effects include microglial activation, increased lipid peroxidation, and neuroinflammation as well as impaired adult neurogenesis. In most cases, the effects of diesel exhaust were more pronounced in male mice, possibly because of lower antioxidant abilities due to lower expression of paraoxonase 2 [60].

While some studies have noted a link between maternal smoking and autism, or in autism subgroups [18,61–63], others have not and a large meta-analysis has concluded no effect [64]. However, the enrichment of ASG’s applies to individual components (some but not all) of cigarette smoke, rather than to overall smoking. In addition, given the gene/environment interaction, it is likely that smoking risk might also be gene-dependent and only seen in susceptible individuals. Meta-analysis based solely on environmental factors does not take this into account. Given the focus of many chemicals on subsets of genes this likely applies to many meta-analysis studies, whether genetic or environmental. This also provides a plausible explanation for the many divergent results in genetic and epidemiological studies.

**Figure 7.**
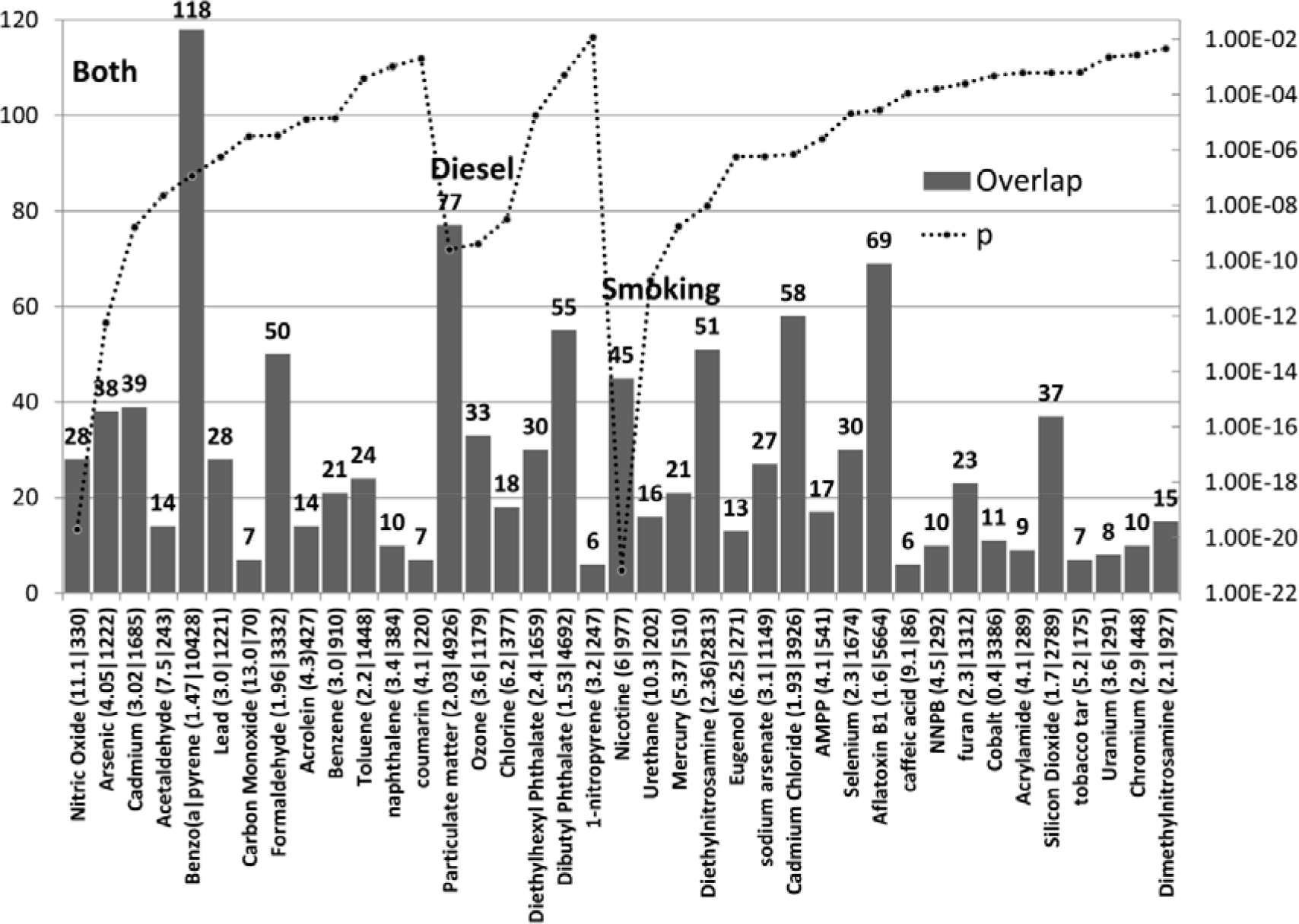
The number of ASG’s affected by compounds found in cigarette smoke or diesel exhaust or in both. N ASG’s =lef axis, p values =right axis). The enrichment ratio and the total number of genes affected by each compound are shown after each compound name.

### Endogenous compounds targeting the autism genes

Neurotransmitters, hormones and key endogenous signalling and other metabolites are the agents through which genes and environmental factors act to influence pathology and behaviour. By inference, pollutants that target the same genes/proteins as those related to endogenous messengers must interfere with their function. As shown below, many of the endogenous agents that target autism genes clearly relate to autism pathology and behaviour.

### Autism genes are targeted by relevant hormones and transmitters (Fig 8)

Compounds with the most significant enrichment scores (tretinoin (=all-trans retinoic acid), melatonin, progesterone and estradiol) and other hormones (thyroxine, triiodothyronine, corticosteroids, calcitriol (1,25-dihydroxyvitamin D3, the hormonally active metabolite of vitamin D)) demonstrate a key influence of retinoids, sex hormones and steroids that is relevant to the suspected role of environmental endocrine disruptors in autism [16, 65]. Lower nocturnal melatonin or melatonin metabolite concentrations have been observed in autism and melatonin treatment improves sleep quality and daytime behaviour in autistic subjects [66, 67].Low vitamin D status during pregnancy or childhood has also been associated with autism [68]. Severe maternal hypothyroxinaemia during early pregnancy has also been linked to an increased incidence of autism in the offspring [69].

The highest scoring neurotransmitters were serotonin, dopamine and noradrenaline, which is generally consistent with current views on the import of these agents in autism pathology and symptomatology [67,70,71]. Sphingosine-1-phosphate (S1P) plays an important role in oligodendrocytes and in myelination [72]. Aberrant myelination, greater than expected for their age in left and right medial frontal cortex and less than expected in the left temporoparietal junction has been noted in autistic children and high serum levels of S1P have been reported in a metabolomics study of autistic subjects [73].As recently reviewed, oxytocin has both beneficial and deleterious effects in autism. While its use to induce labour has been linked to the subsequent development of autism in the children, it can also help in relation to the social skills in autistic patients [74]. Oxytocin reinforces parental and social bonding, and increases anandamide signalling in the mouse nucleus accumbens, an effect that drives social reward [75].

Also of interest is an endogenous aryl hydrocarbon receptor (AHR) ligand ( 2-(1′H-indole-3′-carbonyl)-thiazole-4-carboxylic acid methyl ester (ITE). [76]. AHR is a xenobiotic sensor and the target of dioxins, persistent organic pollutants, polycyclic aromatic hydrocarbons, polychlorinated biphenyls, organochlorine pesticides and endocrine disruptors [77], many of which are the top chemicals targeting the autism genes[78–81]. There appear to have been no studies relating AHR to autism.

Calcium is directly relevant to 3 calcium channels **CACNA1C, CACNA1G, CACNA1H** in the autism gene set. Voltage sensitive calcium channels play an important role in neural function. They are also expressed in the placenta and trophoblast and play an important role in the delivery of calcium to the foetus [82, 83]. Heavy metal cations, particularly lead and mercury, are potent calcium channel blockers but can also permeate these channels, gaining access to the cell [84].

**Figure 8.**
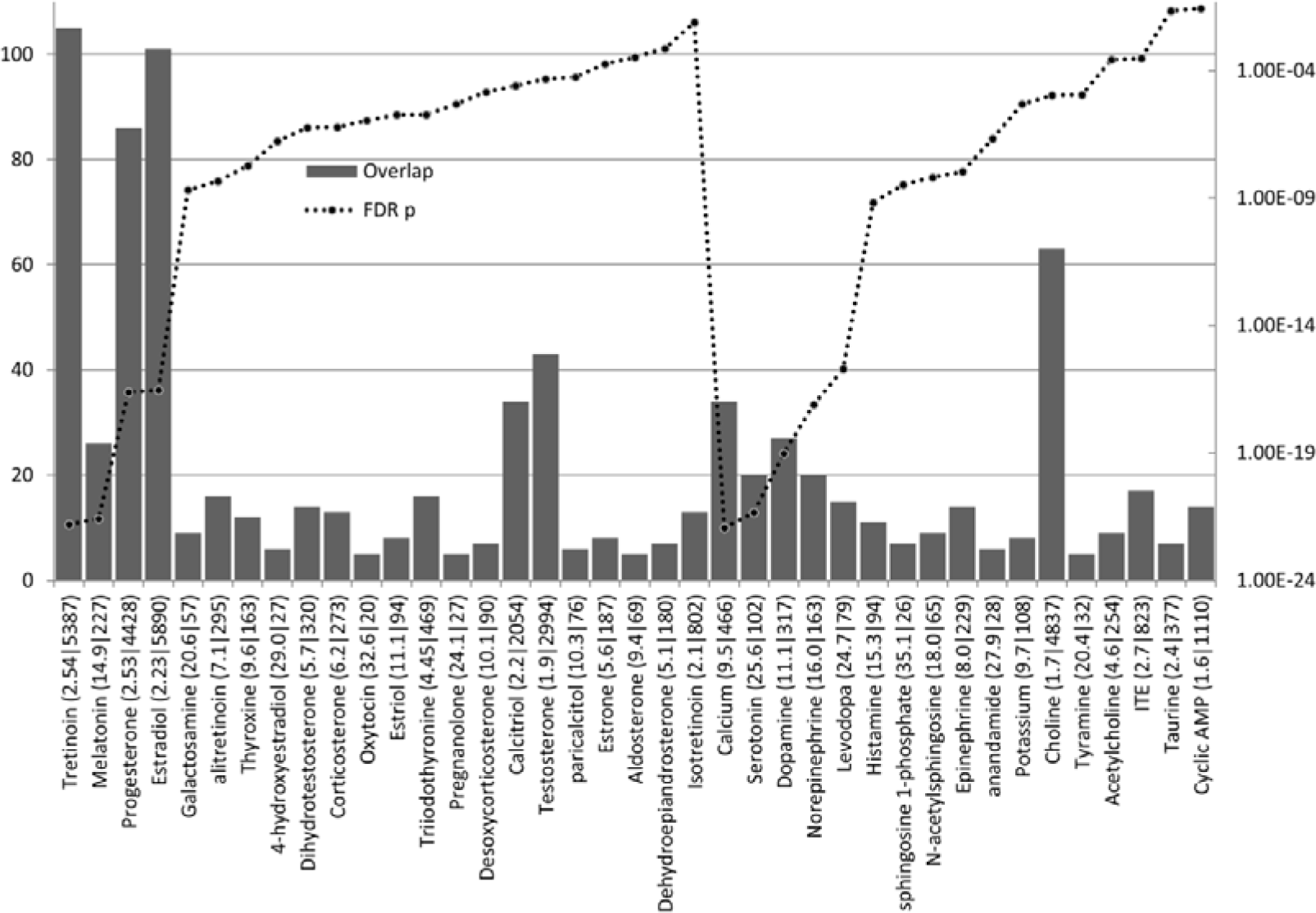
The number of ASG’s affected by Hormones (first batch) and transmitters (second batch: including cations and second messengers). (N ASG’s =left axis, p values =right axis). The enrichment ratio and the total number of genes affected by each compound are shown after each compound name. (ITE = 2-(1′H-indole-3′-carbonyl)-thiazole-4-carboxylic acid methyl ester. Galactosamine is included as it is a constituent of some glycoprotein hormones (follicle-stimulating hormone and luteinizing hormone).

Other endogenous compounds targeting autism genes

These are grouped by general function in Fig 9. They include compounds related to oxidative stress and folate/methionine/homocysteine metabolism which play key roles in autism [85–88] as do cholesterol and fatty acid metabolism [89–92] or inflammation [93–96]․.

Several bile related compounds appear in this figure. Bile acids act as nutrient signalling hormones and activate a number of nuclear receptors including the farnesoid X receptor (NR1H4) the pregnane X receptor(NR1I2), and the Vitamin D receptor. They also activate G-protein coupled receptors including a specific bile acid receptor GPBAR1 which regulates intestinal barrier structure via modification of epithelial tight junctions [97], and a sphingosine-1 phosphate receptor 2 (S1PR2) that has recently been identified as a Nogo-A receptor regulating myelin dependent synaptic plasticity [98] The pregnane X receptor (NR1I2) forms a dimer with a 9-cis retinoic acid receptor [99] and is also activated by Bisphenol A, phthalates and organochlorine pesticides [100] [101] and by many estrogenic and antiestrogenic compounds, both endogenous and environmental [102]. No studies relating bile acids to autism were found in Pubmed, but this area appears to be of interest.

It is important to note that compounds generally considered as beneficial in relation to autism also target autism genes (see discussion caveats). These include folic acid (see above), glutathione and its precursor acetylcysteine which has reported benefits in the treatment of autism and related psychiatric disorders [103] as might vitamins [104, 105].

Thioctic acid or alpha-lipoic acid is an essential coenzyme for α-ketoglutarate and pyruvate dehydrogenase and thus an obligate requirement for energy production[106]. Lipoic acid is able to protect against the effects of Bisphenol A or Bi-n-butyl phthalate on testicular mitochondrial toxicity [107, 108]. In various other models it also protects against the toxic effects of acetaminophen [109], acrolein [110], cyclosporine [111], indomethacin [112], paraquat [113] and rotenone [114] as well as cypermethrin [115], dimethoate, glyphosate and zineb [116],chrysene [117] lindane [118] and Tetrachlorodibenzodioxin [119]. Lipoic acid and other antioxidants have also been used in the clinical management and prevention of heavy metal intoxication [120]. The targeting of autism genes by this product may thus reflect beneficial rather than deleterious effects and, in particular, lipoic acid protects against a large number of toxicants that target autism genes and which have been implicated in the disorder. It has not been analysed in epidemiological studies or tested in the clinic, and blood or tissue levels do not appear to have been measured in pregnancy, neonates or autistic children.

The effects of some fatty acids and carnitine are also oriented towards the ASG’s. Faecal levels of acetic, butyric, other short chain fatty acids and ammonia are increased in autistic children, related to microbiome alterations [121, 122]. Reduced serum carnitine and linoleic acid levels and modified omega3/omega6 fatty acid ratios have also been noted in autism [123].

**Figure 9.**
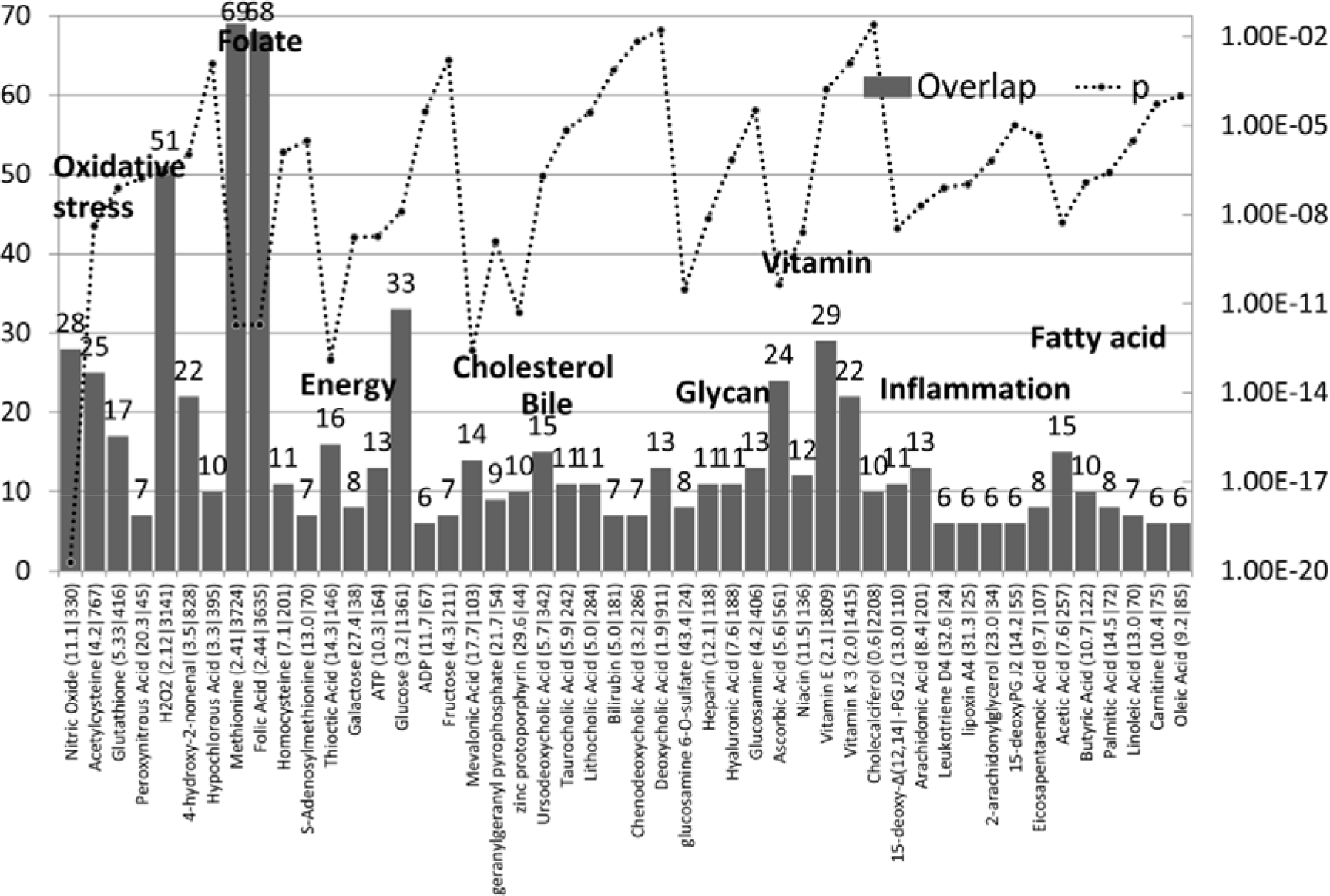
The number of ASG’s affected by diverse endogenous compounds (N ASG’s =left axis, p values =right axis). The enrichment ratio and the total number of genes affected by each compound are shown after each compound name. The compounds are organised in relation to their general function.

### Food additives and compounds in cosmetics targeting the autism genes (Fig 10)

70 compounds in the EAFUS list >5 autism genes (P<0.05). The interactomes of several compounds which might be considered beneficial (folic acid, methionine, ascorbic acid, niacin and oleic acid) were significantly enriched in autism genes. (Methionine (69/3724: 2.41 fold: P= 2.E-12); Folic Acid (68/3635: 2.44 fold: P=2.03E-12); Ascorbic Acid (24/561: 5.6 fold: P=4.52E-11)

Niacin (12/136: 11.5 fold: P=2.58E-09); Oleic Acid (6/85: 9.2 fold: P=0.0001).

Ammonium chloride, the highest scoring compound, derived from burning coal is also used as a flavour enhancer. Ammonium hydroxide in brine solutions is used as a meat tenderiser (363) and is also widely used in food processing to increase pH, while ammonia gas is used to kill bacteria in ground beef (364). NH4Cl might be considered as a potential by-product of such procedures due to reaction with salt or gastric hydrochloric acid. No reports in relation to autism could be found. NH4Cl (Fig 11) increases the permeability of pial venular capillaries to Lucifer Yellow, as does histamine (Fig 10) [124]. It is not practical, given space limitations, to discuss all of these compounds whose relationships with autism or to barrier function remain to be analysed. There are several however that are perhaps of more interest than others due to their extensive use (aspartame, a constituent of over 6000 food products) or as anticaking agents that are also constituents of widely used sunscreens (Titanium dioxide, silicon dioxide, and zinc oxide).

Aspartame acts via sweet taste receptors TAS1R2 /TAS1R3 [125] and also activates transient receptor potential heat and inflammation sensitive channels (TRPV1). These are involved in metallic taste perception as they are also activated by copper, zinc and iron sulphates [126]. **TAS2R1**, within the autism gene set, is a bitter taste receptor. Recent evidence suggests that such receptors, also found in areas outside the mouth, may activate defensive mechanisms against noxious chemicals including cytokine and immune systems. In the human lung, TAS2 receptors are expressed in the cilia that sweep harmful chemicals, particles, and microbes from the airways [127]. TAS2 receptor activation in nasal cells results in the secretion of antimicrobial peptides, an effect inhibited by TAS1R2 /TAS1R3 sweet activation [128]. Thus, aspartame, excessive glucose and other sweet substances activating TAS1 receptors would be expected to inhibit the clearing of pathogens and noxious chemicals stimulated by TAS2 receptor activation.

Microbiome profiling has shown that low-dose aspartame, which has also been implicated in the development of obesity and metabolic disease, increases total bacteria, the abundance of Enterobacteriaceae and Clostridium leptum in diet-induced obese rats. It also increases the serum levels of the short chain fatty acid propionate [129].High levels of faecal enterobacteria and Clostridial families have also been reported in autism [130]. The intracerebroventricular administration of propionate in rats induces behavioural and pathological signs that are relevant to autism [131–134].

Titanium and silicon dioxide (silicon dioxide (37/2789 1.7 fold P=0.0006: not on figure) are used, often in nanoparticle form, in a large number and variety of commercial products including pigment colours, anti-bacterial and other pharmaceutical components, ultraviolet radiation scavengers (sunscreens), as well as in cosmetics. Both are also food additives used as anticaking agents or colorants [135, 136]. Their risks are generally uncharacterised in epidemiological studies although they are manufactured and used worldwide in large quantities [137]. Zinc oxide is also used as a sunscreen.

Titanium dioxide nanoparticles are internalised by human neuronal SHSY5Y cells and induce dose-dependent cell cycle alterations, apoptosis, and genotoxic effects that appear to be unrelated to oxidative damage [138]. In contrast, in human epidermal cells, they reduce glutathione and increase lipid hydroperoxide and reactive oxygen species levels, leading to genotoxicity via oxidative DNA damage [139]. Titanium nanoparticles also suppress angiogenesis [140, 141].

Both titanium and silicon dioxide nanoparticles cross the placental barrier in mice and can be found in foetal liver and brain following maternal administration. Such treatment results in smaller uteri and foetuses [142]. Titanium dioxide nanoparticles accessing the nasal or pulmonary route are also translocated to the brain or the systemic circulation and thence to other organs [143]. The prenatal administration of titanium dioxide nanoparticles in rats results in oxidative damage to brain nucleic acids and lipids in neonates and a depressive behavioural spectrum in adulthood [144]. It also increases frontal cortical and neostriatal dopamine levels in the offspring [145]. In mice, prenatally administered titanium dioxide increases allergic susceptibility; impairs spermatogenesis and induces marked gene expression changes in the brains of the offspring [146]. Prenatally exposed mice subsequently display signs of lung inflammation and increased susceptibility to asthma and avoid the central zone of the open field, while the female offspring show enhanced prepulse inhibition [147, 148]. Both silicon and titanium dioxide nanoparticles activate inflammatory cascades in microglia and the supernatants collected from the treated microglia are cytotoxic to PC12 neuronal cells [149]. Titanium dioxide nanoparticles are internalised by microglial cells. They inhibit cell adhesion and increase cytoplasmic membrane permeability to propidium iodide associated with a loss of mitochondrial transmembrane potential and an overproduction of superoxide [150]. Titanium dioxide nanoparticles adsorb bisphenol A and this complex is taken up by zebrafish causing a reduction in the plasma concentrations of estradiol (E2), testosterone, follicle-stimulating hormone, and luteinizing hormone [151].

Beta-ionone (EAFUS/cosmetics) is a flower carotenoid derivative contributing to the scent of roses. It is also formed in animals by beta-carotene oxygenase 2 (BCO2) which converts betacarotene to β-10′-apocarotenal and β-ionone, en route to the synthesis of Vitamin A [152]. It does not activate retinoid receptors RARA or RARB [153] but binds to a retinol binding protein, beta-lactoglobulin B, involved in the oral delivery of retinol to neonates [154]. It is a potent inhibitor of a mouse retinal dehydrogenase (raldh4: human homologene = ALDH8A1). These enzymes catalyze the dehydrogenation of retinal into retinoic acids, which are required for embryogenesis and tissue differentiation [155].

Tetramethylpyrazine/ligustrazine is a Chinese herbal ingredient, that in combination with other herbal extracts, Ferulic acid (7/54: 16.9 fold: P=5.82E-07) and tetrahydropalmatine (3/150: 2.6 fold: P=0.08 (NS))inhibit the growth of ectopic endometrial tissue in endometriosis rats. This preparation also decreased the levels of estradiol and suppressed the expression of GnRH, FSH and LH, as well as decreasing estrogen receptor expression [156].

**Figure 10.**
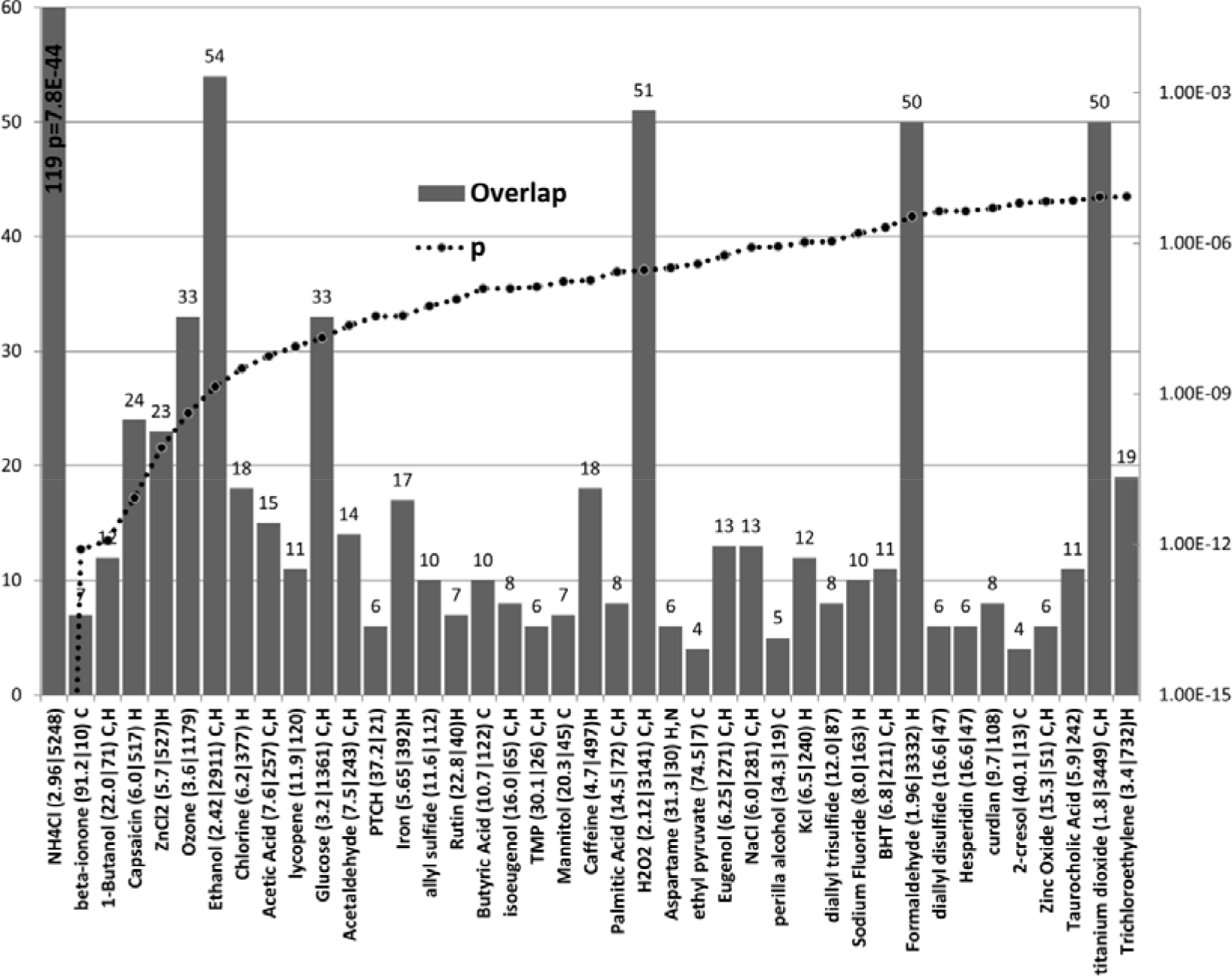
Compound on the EAFUS list that target the ASG’s The maximum left and minimum Y axes are truncated for clarity (NH4Cl affected 119 autism genes: p= 7.78E-44). BHT = butylated hydroxytoluene; PTCH = protocatechuic acid (a major metabolite of antioxidant polyphenols found in green tea.) TMP= tetramethylpyrazine. Compounds also found in cosmetics are appended with C and those in household products with H.

### Cosmetic ingredients targeting autism genes

Several of these are also in the EAFUS list (see above) and may be used in both as solvents or fragrances and only those specific to cosmetics or not dealt with above are shown in Fig 11.

Tretinoin, (all-trans retinoic acid) is the highest scoring compound. It is used for acne and as an anti-ageing component in face creams [157] and is available, without prescription, on many websites. Tretinoin treatment in pregnant rats results in postnatal mitochondrial complex 1 dysfunction in the cerebellum of the offspring [158] and has also been shown to increase levels of fear and anxiety in offspring [159]. Gestational treatment also results in a delayed appearance of the cerebellar righting reflex and reduces open-field activity in the offspring. In addition the offspring show impaired motor coordination and motor learning ability coupled with a reduction in the cerebellar size and impairment in the cerebellar foliation profile [160, 161]. A 3 day exposure to 2.5 mg/kg tretinoin (gestational days 11-13) produces a 10% reduction in weight of cerebellum at 4 weeks of age, not accompanied by other malformations [162]. In rats treated with retinoic acid at gestational day E10, the foetuses show structural changes similar to humans with Arnold-Chiari malformation, including downward displacement of the cerebellum to just above the foramen magnum and compression of the developing medulla into a small posterior fossa[163]. A recent MRI study has commented on the co-existence of Chiari malformation with some paediatric autism patients [164]. The targeting of the cerebellum by tretinoin is particularly relevant given that cerebellar abnormalities are a consistent feature of autism [165–167]. The transfer of retinoic acid across pig skin is increased by exposure to particulate matter containing polycyclic aromatic hydrocarbons [168].

Acetovanillone inhibits the free radical superoxide generator NADPH oxidase [169]. The activity of this enzyme is decreased in granulocytes and lymphocytes of autistic children contributing to a spectrum of mitochondrial malfunction in these cases [170, 171].

Patchouli alcohol is used in a number of perfumes. It is endowed with a variety of potential benefits including anti-oxidant and inflammatory effects [172], inhibition of indomethacin and ethanol-induced gastric ulcers [173] as well as anti-bacterial and antiviral effects [174, 175]. However it also decreases cell growth in MCF7, BxPC3, PC3, and HUVEC cells and downregulates histone deacetylase HDAC2 in human colorectal cancer cells [176]. HDAC2 is a valproate target also forming a complex with the Rett syndrome gene **MECP2** [177, 178].

Limonene is an inhibitor of protein farnesyl transferase (FNTA FNTB) and protein geranylgeranyl transferase (PGGT1B) through which it exerts anticancer properties [179]. Farnesylation, a lipid posttranslational modification, adds farnesyl diphosphate to cysteine residues allowing substrates to interact with membranes and protein targets. It is essential for embryonic development [180]. Farnesyl and geranylphosphate play a role in angiogenesis in human umbilical endothelial cells [181, 182]. Limonene is metabolised by CYP2C9 and CYP2C19[183] both of which metabolise progesterone, while testosterone is a substrate for CYP2C19 [184].

Nonylphenol is a persistent endocrine disruptor used in home maintenance products that is also ubiquitous in foodstuffs for babies and toddlers commercially available in Germany [185] and in many other foods including human breast milk in Europe [186, 187]. Urinary Nonylphenol levels have been associated with spontaneous abortions [188], and plasma levels with precocious puberty (also observed with mono(2-ethylhexyl) phthalate, t-octylphenol, daidzein, equol, and genistein)[189]. Nonylphenol and other compounds including dioxins, polychlorinated biphenyls, organochlorine pesticides, bisphenol A, and phytoestrogens have also been detected in umbilical cords and cord sera in Japan [190].

Glycyrrhizic acid is the sweet ingredient of the liquorice root. It lowers the activity of 17 beta-hydroxysteroid dehydrogenase, which converts androstenedione to testosterone and decreases serum testosterone levels in women, and ovarian testosterone levels in rats [191, 192].

**Figure 11.**
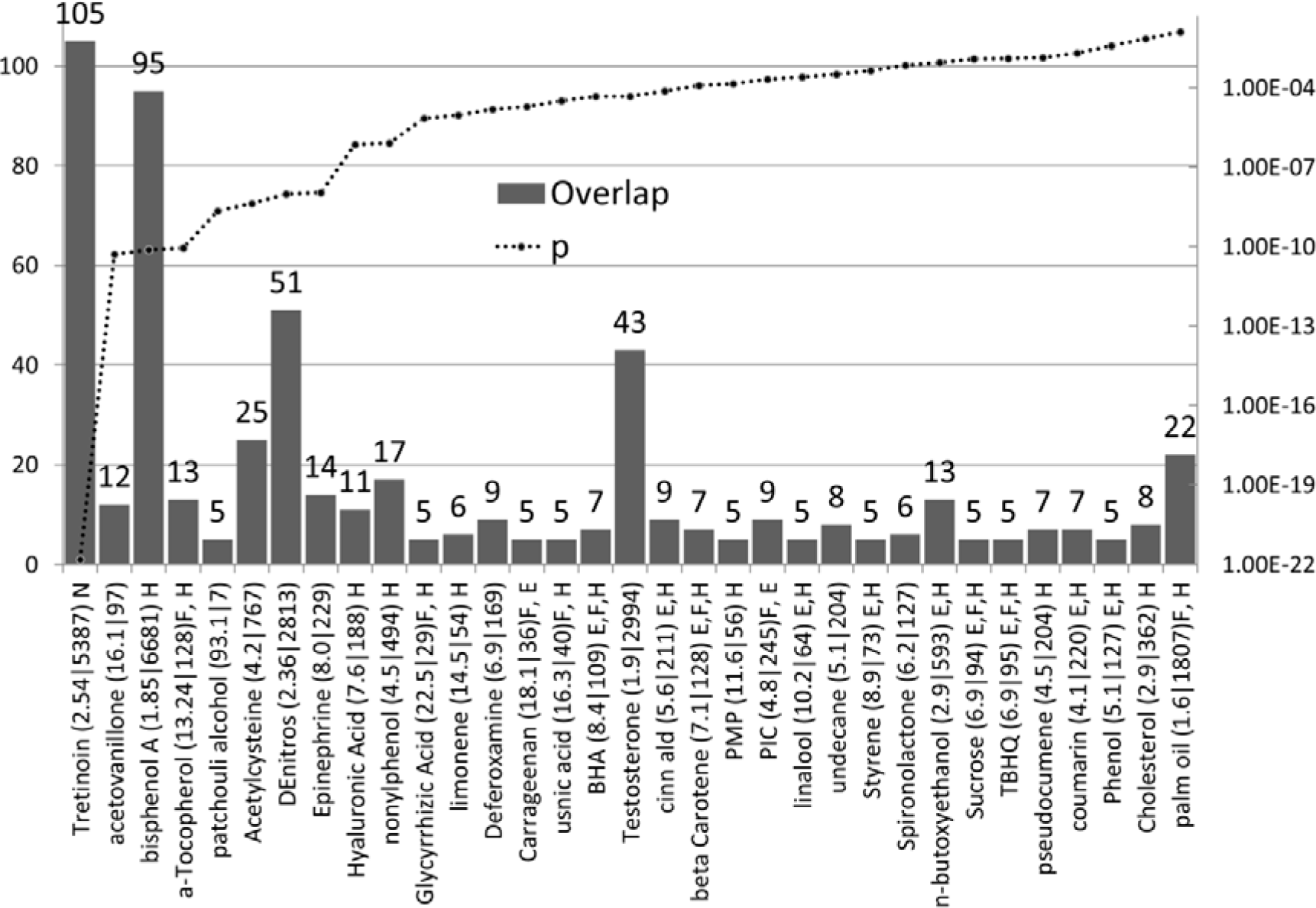
The number of ASG’s affected by compounds in cosmetics (N ASG’s =left axis, p values =right axis). The enrichment ratio and the total number of genes affected by each compound are shown after each compound name. The presence of these compounds in food (F), the EAFUS list (E), nutraceuticals (N) or household products (H) is also indicated. a-Tocopherol = alpha-Tocopherol; BHA= Butylated hydroxyanisole; DeNitros = Diethylnitrosamine; PMP= phenylmethylpyrazolone; PIC= phenethyl isothiocyanate; TBHQ = 2-tert-butylhydroquinone.

### Compounds affecting barriers or respiratory cilia

As previously reported[38], many of the autism genes in this set are involved in barrier functions across several different boundaries (blood/brain, skin, intestinal and placental) and also in the control of respiratory cilia that clear the airways of noxious particles. Evidently, environmental chemicals have to traverse such boundaries. In addition, some also have deleterious effects on barrier or cilia function.

Several pesticides (malathion and lead acetate, Chlorpyrifos or a combination of the insect repellent, DEET (N,N-Diethyl-meta-toluamide) and permethrin) are able to disrupt the blood brain barrier in animal models [193] and nicotine and smoking disrupt brain microvasculature and the blood brain barrier[194]. Long-term air pollution in cities relates to neuroinflammation, an altered innate immune response, disruption of the blood-brain barrier, ultrafine particulate deposition, and accumulation of beta-amyloid in children and young adults [195]. Air pollution also disrupts epithelial and endothelial barriers and triggers autoimmune responses involving tight junction and neural autoantibodies [196].

Nanoparticles from aluminium, silver or copper increase spinal cord pathology after trauma, an effect correlated with breakdown of the blood-spinal cord barrier [197]. NH4Cl increases the permeability of pial venular capillaries to Lucifer Yellow, as does histamine.[124]. The transfer of retinoic acid across pig skin is increased by exposure to particulate matter containing polycyclic aromatic hydrocarbons [168].

With regard to respiratory cilia, cigarette smoke decreases beat frequency and cilia length is reduced in healthy smokers. Long-term exposure to cigarette smoke leads to reduced numbers of ciliated cells in mice [198, 199]. A combination of cigarette smoke and alcohol also decreases ciliary beat frequency in bovine primary ciliated bronchial epithelial cells [200]. Chlorocresol, a disinfectant, decreases ciliary beat frequency in human nasal epithelial cells [201], and the insecticide deltamethrin provokes respiratory ciliary damage in rats [202]. The fungicide benomyl and its metabolites, butyl isocyanate and carbendazim, decrease ciliary beat frequency in canine tracheal epithelial tissue [203]. Progesterone inhibits cilia beat frequency in human lung and cultured primary human airway epithelial cells, an effect inhibited by 17beta-estradiol [204]. No effects could be found in relation to endocrine disruptors, although they might be expected to exert effects in relation to those of these steroid hormones. Ciliary function is also compromised by vanadium, vanadium-rich oil-fired fly ash and cadmium [205, 206]. As noted above, bitter taste receptors increase cilia function, and these are inhibited by sweet taste receptors activated by aspartame and glucose.

Such deleterious effects are likely to modify the intake of many other compounds.

### Ecological pollution and bioaccumulation

Many compounds used in cosmetics or as food additives can be directly absorbed or ingested and pesticide sprays and volatile compounds inhaled. While the concentrations of some may well be too low to elicit direct toxicity individually, a further problem relates to the disposal of multiple products down drains or in waste dumps from where they can seep into the air and water tables. Such compounds can be concentrated by the food web (bioaccumulation). In a Mediterranean river diclofenac, gemfibrozil, and the flame retardant TBEP were detected in water, biofilm and in macroinvertebrate taxa [207]. Detectable levels of polychlorinated biphenyls and pesticides, with evidence of thyroid dysfunction have been reported in wild fish in the San Francisco bay area [208]. Compounds in pesticide sprays can also travel long distances. Nonylphenol polyethoxylate enters the environment as an inert ingredient in pesticide sprays, which are used extensively in California’s Central Valley. In the distant Eastern Sierra Nevada Mountains, trace amounts of 4-nonylphenol can be found in surface water, snow, and atmospheric deposition, some with yearly average concentrations as high as 9 μg/L [209]. Such compounds exist in multiple permutations in relation to environmental contamination. The effects of the various compounds, as illustrated in the figures above, apply to individual compounds, but the real life situation involves multiple ingredients in food or cosmetics and diverse mixtures of environmental pollutants with additive effects. The enrichment of autism genes in the effects of these compounds must therefore be viewed in this context. In an American study in 2011, certain polychlorinated biphenyls, organochlorine pesticides, perfluorinated compounds, phenols, polybrominated diphenyl ethers (flame retardants), phthalates, polycyclic aromatic hydrocarbons, and perchlorate were detected in 99-100% of pregnant women [210].This ecological problem applies to agrochemical and industrial pollutants and likely to hundreds of biologically active compounds in food, cosmetic, drug and household products. Concerns relating to these products in relation to autism and other neurodevelopmental and cognitive disorders has recently been raised in The TENDR Consensus Statement, a call to action to reduce exposures to toxic chemicals [211].

**Caveats:** There are numerous caveats. Firstly, this is a comparison of two lists of gene symbols, with no indication of weight (relative importance in relation to specific genes or processes) or directionality (i.e. does the compound activate or inhibit, or are the effects on binding, transcription or phosphorylation, etc), although these can be found within CTD and in the literature for any interaction of interest. This type of enrichment applies to toxicant chemicals, but also to those that might be beneficial (e.g. folic acid, lipoic acid, or glutathione), or a mixture of both (e.g. Oxytocin).In many cases, for example pesticides, heavy metals, bisphenol A, phthalates, valproate, etc.), a link to autism is supported by epidemiology and/or by animal studies in relation to development. Related compounds not yet studied in autism, particularly atrazine and other pesticides, blue asbestos or known endocrine disruptors can hardly be considered as benign. Other compounds, for example aspartame, titanium dioxide or sodium fluoride possess endocrine disrupting or other toxic effects relevant to neurodevelopment. As stated on the CTD website, such data can be used for hypothesis testing. It is impossible to predict whether any uncharacterised compound plays a causal role in autism, but these data can at least provide a long list worthy of further investigation in epidemiological and animal studies.

## Discussion

The specific question posed by this type of analysis is not whether any compound affects autism genes/proteins, but whether it affects more autism genes than would be expected from the overall toxicological profile of that compound. If such is the case, one might assume that there is a particular relationship between genes and environment that suggests that the genetic polymorphisms, as well as disrupting key autism pathways related to pathology, also affect the ability of certain toxicants to exert their effects via the same genes or proteins. One might therefore expect that many of these genes, also related to barrier function, modify the absorption, metabolism, excretion or physiological effects of the toxicants. In several cases, this has been shown to be the case, and certain autism polymorphisms do affect these parameters[3, 38], although this has not been tested for all of the many genes or chemicals involved.

In relation to these questions, several hundred compounds selectively target multiple members of this particular group of 206 genes. 6338 unidentified compounds in CTD did not affect any autism gene, while the effects of others were not significant, showing a degree of specificity. Within this group of significant compounds are the majority of the compounds suspected to be implicated in autism including pesticides, heavy metals, and industrial pollutants, Bisphenol A and phthalates and several drugs, fluoxetine and other SSRI’s, as well as acetaminophen, valproate and certain drugs used in labour. This exercise also returned all of the general classes of compounds suspected to be implicated in autism, including particulate matter and other components of diesel exhaust, polyhalogenated biphenyls, flame retardants, polycyclic aromatic hydrocarbons, persistent organic pollutants and endocrine disruptors. The endogenous hormones and transmitters targeting these genes are also highly relevant to endocrine disruption and to the key transmitters related to autism (retinoids, sex steroids, thyroxine, melatonin, folate, dopamine, and serotonin) and to the processes implicated in pathology (compounds related to oxidative stress, folate/methionine/homocysteine, inflammation or myelination). Many more compounds were identified, which due to the cumulative nature of many of these exposures, might also play a role. Overall, these data suggest that this type of enrichment analysis reliably identifies key compounds involved in autism, and that it might therefore also have predictive value.

In the present analysis, a large number of compounds, some with clearly relevant toxic effects (e.g. titanium dioxide, tretinoin, or aspartame) can really only provide a list worthy of consideration in epidemiological and toxicological studies, particularly during pregnancy and in relation to neurodevelopment and autism.

Many of these compounds are considered safe by government authorities, but no regulatory toxicological studies could have taken into account the possibility that toxicity might be determined by the same genes that govern susceptibility to autism. This problem could perhaps be addressed using a range of compounds and banked stem cells or tissues from autistic patients or their parents to analyse whether toxicant properties differ in autism cells. A large number of chemicals relate to many autism genes suggesting that the two act in concert and that the rise in the incidence of autism is likely to be chemically driven, in a gene-dependent manner. In this study relating chemicals or environment to genes, it seems that genes and environment are indissociable and that the susceptibility genes themselves may constitute one of the strongest arguments for a causal effect of the environment, as it is towards their products that multiple environmental influences are selectively directed, and via the agency of the gene products that the pathology must be induced, or the toxic products allowed to pass or act.

There appears to be no known reason to suppose that the same genetic variants did not exist in the population prior to the autism epidemic, but a modified environment might have rendered them more relevant to autism. This is akin to the classical population genetics example of the peppered moth. The genes controlling its mottled colouring originally conferred protection from birds, due to camouflage on similarly marked tree bark. Such trees were blackened by industrial soot pollution, and the same genes now conferred a high risk of predation [212], a situation reversed by clean air acts in the UK and USA [213].

The solution to autism prevention may thus similarly reside in the detection, avoidance and removal of the pollutants, a task involving the development of stricter and more appropriate toxicological and environmental controls at governmental level worldwide, as already proposed in the recent TENDR Consensus Statement (Targeting Environmental Neuro-Developmental Risks) [211].

This research did not receive any specific grant from funding agencies in the public, commercial, or not-for-profit sectors.

The authors report no conflicts of interest.

